# The cryo-EM structure of trypanosome 3-methylcrotonyl-CoA carboxylase provides mechanistic and dynamic insights into enzymatic function

**DOI:** 10.1101/2023.11.27.568864

**Authors:** Adrián Plaza-Pegueroles, Inna Aphasizheva, Ruslan Aphasizhev, Carlos Fernández-Tornero, Federico M. Ruiz

## Abstract

3-methylcrotonyl-CoA carboxylase (MCC) catalyzes the two-step, biotin-dependent production of 3-methylglutaconyl-CoA, an essential intermediate in leucine catabolism. Given its critical metabolic role, deficiencies in this enzyme associate with organic aciduria, while its overexpression is linked to tumor development. MCC is a dodecameric enzyme composed of six copies of each α- and β-subunit. We present the cryo-EM structure of the endogenous MCC holoenzyme from *Trypanosoma brucei* in its soluble, non-filamentous state at 2.5 Å resolution. We unambiguously observe the position of biotin, covalently-bound to the BBCP domain of α-subunits and occupying a novel binding pocket next to the active site of a neighboring β-subunit dimer. Moreover, flexibility of key residues at the α/α- and α/β-subunit interfaces enables pivoting of α-subunit trimers to sequentially approach the otherwise distant active sites for the two steps in MCC catalysis. Our results provide a structural framework to understand the enzymatic mechanism of eukaryotic MCCs and assist drug discovery against trypanosome infections.

## Introduction

Biotin-dependent carboxylases are extensively distributed in nature, having a key role in the metabolism of different substrates such as fatty acids, carbohydrates or amino acids (Tong 2013). Four of these enzymes are known in eukaryotes, i.e. acetyl CoA carboxylase (ACC), pyruvate carboxylase (PC), propionyl CoA carboxylase (PCC), and 3-methylcrotonyl-CoA carboxylase (MCC). MCC is responsible for catalyzing the carboxylation of 3-methylcrotonyl-CoA, a key intermediate in the leucine degradation pathway, converting it into 3-methylglutaconyl-CoA. Subsequent steps in the breakdown of this branched-chain amino acid ultimately generate acetyl-CoA and succinyl-CoA, used for energy production or in other metabolic processes (Gondáš et al. 2022; Gallardo et al. 2001). MCC, present in the mitochondrial matrix, is highly expressed in liver and kidney (Obata et al. 2001; Hector et al. 1980).

In humans, deficiencies in MCC can lead to a rare genetic disorder known as 3-methylcrotonyl-CoA carboxylase deficiency or 3-methylcrotonylglycinuria (Baumgartner et al. 2001; Gallardo et al. 2001; Holzinger 2001; Desviat 2003; Baumgartner 2005), which is the most frequently-detected organic aciduria in newborns worldwide (Fonseca et al. 2016). Clinical manifestations of this autosomal recessive defect include hypoglycemia, ketosis, hypotonia, seizures, and developmental and psychomotor delays (Stadler et al. 2006; Grünert et al. 2012; Nguyen et al. 2011; Cheng et al. 2023). On the other hand, increased MCC expression promotes proliferation of several cancer cells, being crucial for the oncogenesis of hepatocellular carcinoma (Chen et al. 2021). Moreover, MCC expression in human astrocytoma, glioblastoma, meningioma and oligodendroglioma tumors indicates the capacity of brain tumor cells to use leucine as a substrate for their metabolism, indicating leucine metabolism as a novel target in cancer treatment (Chen et al. 2021). Augmented MCC expression was also observed in *Leishmania* species resistant to antimony, which suggests MCC as a putative drug target against leishmaniosis (Tasbihi et al. 2020). Additionally, MCC overexpression enhances adaptation of *Arabidopsis thaliana* to alkalized soil, a major problem impacting agricultural production, by increasing tolerance to sodium carbonate and bicarbonate (Wang et al. 2022).

The MCC enzymatic activity involves two subsequent steps (Knowles 1989), taking place at different and distant active sites (Wood and Barden 1977; Tong 2017) (Fig. 1A). The first step is a Mg^2+^/ATP-dependent carboxylation of biotin (vitamin B7 or vitamin H) using bicarbonate as donor (Knowles 1989; Attwood and Wallace 2002). The active center for this reaction resides in MCC α-subunits. The second step is a transfer of the carboxyl group from carboxylated biotin to 3-methylcrotonyl-CoA, which takes place at an active site formed by two MCC β-subunits.

**Figure 1.**
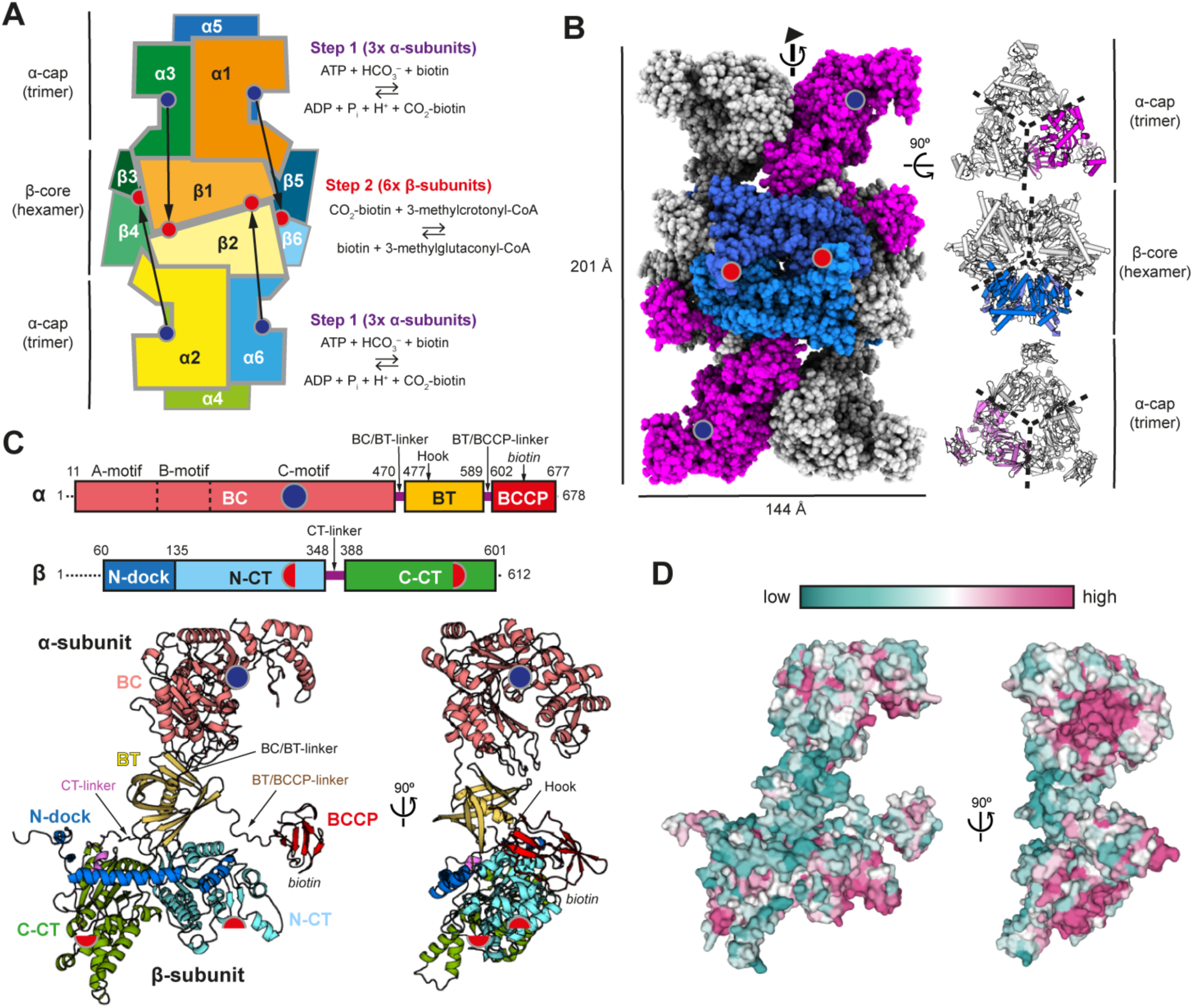
Cryo-EM structure of TbMCC. (A) Schematic representation of dodecameric MCC, showing the hexameric β-core and both trimeric α-caps. Active sites for both steps in the enzymatic reaction are shown with blue and red circles for the first and second steps, respectively. Arrows indicate the necessary movement of BCCP domain for biotin shuttling between active sites. Two arrows, from α5 to β3/β4 and from α4 to β5/β6, are not depicted as they occur at the back of the drawing. (B) Overall structure of *T. brucei* MCC with one of the three twisted αββα segments colored in magenta and blue for α- and β-subunits, respectively. (C) Schematic and ribbon representations of α- and β-subunits of TbMCC, showing domains and motifs in both chains. (D) Degree of conservation of residues in the surface of an α/β dimer.

Current knowledge on the architecture of this enzymatic complex was primarily derived from the crystal structure of MCC from the bacterium *Pseudomonas aeruginosa* (PaMCC) (Huang et al. 2012). The functional complex consists of six copies of each of α- and β-subunits, giving rise to an elongated, dodecameric barrel (Fig. 1A). β-subunits assemble into a hexameric core, that can be considered as a trimer of tail-to-tail β-dimers, positioned at the central part of the barrel. Two α-subunit trimers cap the β-subunit core on opposite sides. α-subunits comprise three domains connected by two short linkers (Hu et al. 2023). The N-terminal biotin carboxylase (BC) domain is placed at the edge of the barrel and contains the α-subunit active site (Fig. 1A, purple circles). The C-terminal biotin carboxyl carrier protein (BCCP) domain contains the Met-Lys-Met biotinylation motif (Samols et al. 1988; Chapman-Smith and Cronan 1999), where the exposed central lysine is the target for covalent binding of biotin to form biocytin (Reche et al. 1998; Athappilly and Hendrickson 1995). The middle BT domain connects BC and BCCP domains via extended linkers and establishes interactions with the adjacent β-subunit. β-subunits are composed of an N-terminal N-dock domain, followed by two carboxyltransferase (N-CT and C-CT) domains that are connected through the CT-linker. The β-subunit active site locates at the β-dimer interface, so that each dimer harbors two active sites (Fig. 1A, red circles). The 3-methylcrotonyl-CoA substrate is sandwiched between the C-CT from one β-subunit and two α-helices in the N-CT from the opposite subunit. The active sites of the two enzymatic reactions locate 80 Å apart (Fig. 1A, black arrows), which requires shuttling of the BCCP domain, facilitated by flexibility of the BT-BCCP linker (Huang et al. 2012). This swinging-domain model has also been proposed for other carboxylases (Perham 2000; St. Maurice et al. 2007).

An equivalent architecture has been found in the electron cryomicroscopy (cryo-EM) structure of MCC from the eukaryote *Leishmania tarentolae* (LtMCC), forming straight filaments composed by four to six dodecameric MCC barrels connected though α-subunit trimers (Hu et al. 2023). In this structure, the BT-BCCP linker is partially fixed through an interaction with the N-dock domain of a neighboring β-subunit, which hampers the BCCP domain to reach the α-subunit active site. Based on this, concomitant swinging of the BCCP and BT domains of the same α-subunit was suggested for LtMCC, in the so-called dual-swinging model. However, LtMCC oligomerization immobilizes BC domains, which are flexible only at the filament edges. Additionally, biotin is not found attached covalently to lysine in the BCCP domain in the filaments. These two features suggest that the LtMCC structure represents an inactive state of this enzyme (Hu et al. 2023).

Here, we applied cryo-EM to study the structure of the endogenous MCC holoenzyme from the parasite *Trypanosoma brucei* (TbMCC). The map exhibits an overall resolution of 2.5 Å while regions of the barrel core are resolved to 2.3 Å resolution, which allowed us to reveal relevant structural details. The resulting structure represents a resting state of the enzyme, where the BCCP domain is weakly-bound to an unexpected position next to a the β-subunit active site. Our cryo-EM analysis also provides insights into the dynamics of TbMCC and enables generation of a mechanistic model for MCC activity.

## Results

### Overall structure

While performing a purification from a *T. brucei* lysate, using cartridges containing agarose coupled with streptavidin, we found a barrel-like structure clearly distinguished in negative-stain grids (Fig. S1). The 3D reconstruction derived from this experiment, combined with mass spectrometry fingerprinting of the bands observed in SDS-PAGE, enabled us to identify TbMCC as the main sample component. Using single-particle cryo-EM, we obtained a map at 2.5 Å resolution that allowed full model building of this dodecameric structure (Fig. S2; Table S1). TbMCC presents a barrel-like structure with a height of 201 Å and a diameter of 144 Å (Fig. 1B, left), closely resembling those of PaMCC (Huang et al. 2012) and LtMCC (Hu et al. 2023). The barrel core is formed by a trimer of β-subunits dimers, while two trimers of α-subunits occupy the top and bottom of the barrel. The structure exhibits 3-fold symmetry along the cylinder axis, such that the repeating unit is a twisted segment of αββα where α- and β-subunits form a 60° angle (Fig. 1B, right). Pseudo-twofold symmetry is also apparent between barrel halves but canonical symmetry is disrupted by differential configuration of BC and BCCP domains of α-subunits on each barrel half (Fig. S2). While these domains show poor density in all cases, their density is better defined on one barrel half, which may indicate allosterism. The β-subunit hexamer defines a large cavity (41481 Å^3^) that is capped at the two α/β interfaces (Fig. S3). Two types of tiny pores are formed by positively charged residues, one at the center of each β-subunit dimer and the second between two dimers. While the internal dimensions are small (about 4 Å), minor conformational changes may allow access of small ions into the β-subunit cavity. In addition, each α-subunit trimer defines a cavity (31500 Å^3^) that is open both at the edge of the barrel and through three additional apertures at the α/β interface (Fig. S3).

### α-subunits present a semi-open BC domain and a structured BT-BCCP linker

The N-terminal BC domains of α-subunits (residues 11-470) define the edges of the dodecameric barrel. This domain can be divided in three structural motifs (Fig. 1C-D), as described for a homologous BC enzyme from *Escherichia coli* (Thoden et al. 2000). The A-motif (residues 11-139) and C-motif (residues 216-470) pack against each other at the center of the α-subunit trimer with the A-motif contacting the BT domain of a nearby α-subunit. The middle B-motif (residues 140-215) or ‘BC lid’ extends away from the barrel in a semi-open conformation, as seen when compared with ATP-bound or Apo *E. coli* BC enzyme (Fig. 2A-B). This motif exhibits flexibility, as suggested by poor definition of the map in this region (Fig. S2). Flexibility of the BC lid plays a role in closing the BC active site for catalysis, such that BCCP binding immobilizes the lid (Chou et al. 2009). The C-motif harbors the active site for carboxylation of BCCP-bound biotin. Consistently, residues in this region are highly conserved (Fig. 1D; Fig. S4). While substrates for the first catalytic step are absent in the TbMCC structure, an extra density is observed in a pocket defined by positively-charged residues K247, R303 and R354 (Fig. 2C). Because we used ammonium sulphate during purification, we modeled a sulphate ion in this density that corresponds to the bicarbonate binding site, as seen in the BC site of the PC enzyme (López-Alonso et al. 2022) (Fig. 2D). Comparison with PC suggests that R303 and E307 in TbMCC likely promote bicarbonate deprotonation while R354 stabilizes deprotonated biotin, which is the final CO_2_ acceptor. Interestingly, R354 in TbMCC contacts E250 through a salt bridge that in PC is formed after ATP hydrolysis to prevent the reverse reaction (López-Alonso et al. 2022).

**Figure 2.**
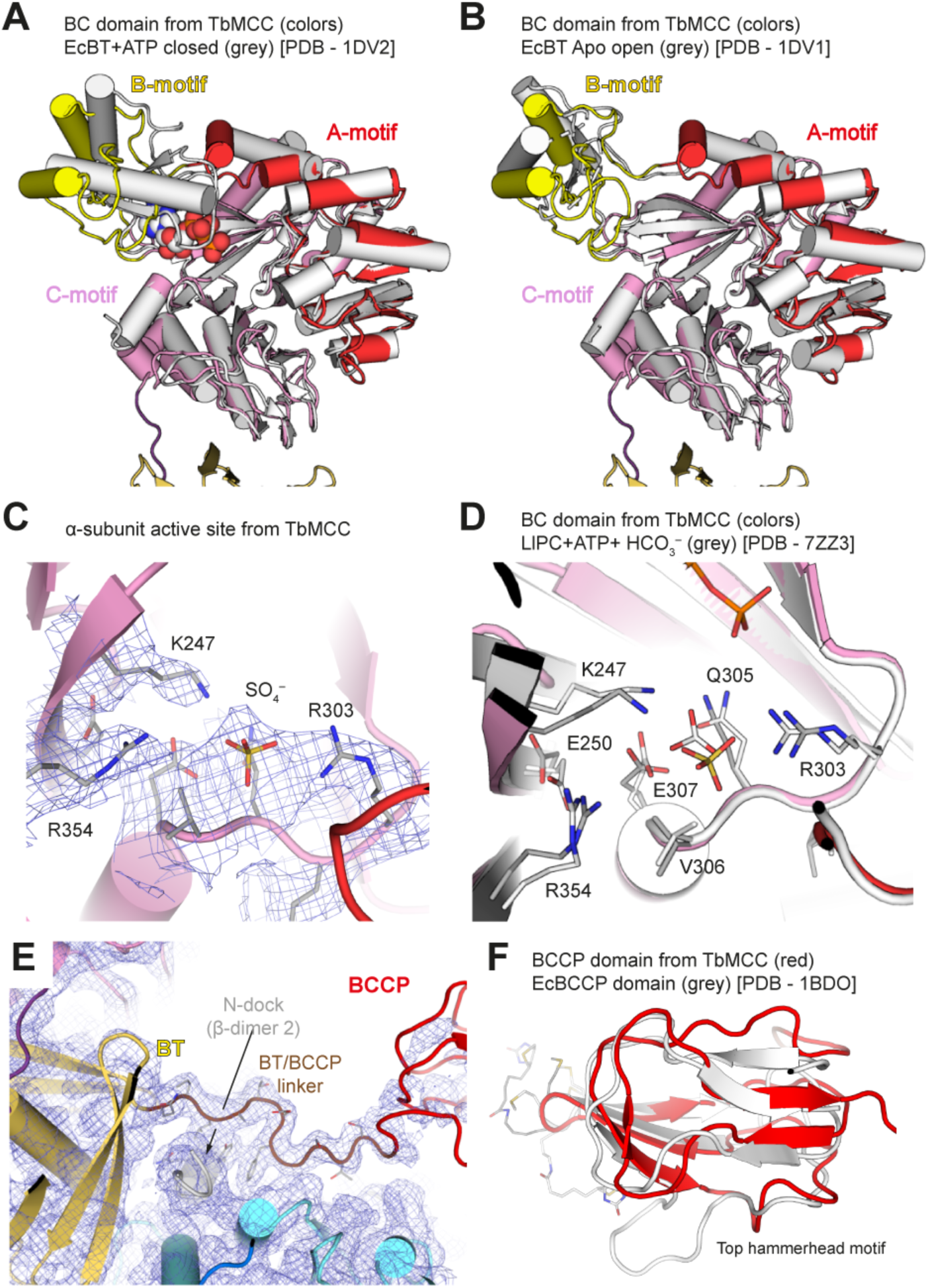
Structural details of α-subunits. (A, B) Structural comparison of the BC domain with biotin carboxylase from *E. coli* complexed with ATP (panel A, PDB - 1DV2) or in the apo form (panel B, PDB – 1DV1), both in grey. The A-, B- and C-motives in the BC domain of TbMCC are shown in red, yellow and pink, respectively. (C) Cryo-EM map showing the bicarbonate binding site where a sulphate ion was modeled. (D) Structural comparison of the BC active site of TbMCC, in pink, with the BC active site in pyruvate carboxylate from *Lactococcus lactis*, in grey (PDB – 7ZZ3), showing the high degree of conservation in sequence and geometry of this region. (E) Cryo-EM map around the BT/BCCP-linker and surrounding regions. (F) Structural comparison of BCCP domain of TbMCC, in red, with a BCCP domain from *E. coli*, in grey (PDB - 1BDO).

The central BT domains of α-subunits (residues 477-589), comprising an axial α-helix enclosed by an eight-stranded β-sheet, locate between BC domains and the β-subunits hexamer (Fig. 1C). Residues connecting the α-helix and the β-sheet in the BT domain form the hook (residues 491-505), which is the major region interacting with the adjacent β-subunit. Nevertheless, this region presents the lowest conservation degree among α-subunits (Fig. 1D; Fig. S4). The BT-BCCP linker (residues 590-601), which is clearly seen in the electron density map (Fig. 2E), interacts with a β-subunit from a neighboring dimer. The BCCP domain (residues 602-677) is placed next to the active site of this β-subunit dimer (Fig. 1B). This domain adopts the capped β-sandwich fold (Athappilly and Hendrickson 1995; Roberts et al. 1999), with β-strands covering two sides of the sandwich and two symmetric hammerhead motives (residues 620-634 and 657-671) (Fig. 2F). This domain exhibits flexibility according to poor definition of the cryo-EM map in this region (Fig. S2).

### β-subunits form active sites that are accessible to the substrates

The N-dock domain (residues 60-135) of β-subunits contains two α-helices (residues 90-115 and 119-130) that extend at the outer rim of the barrel core and establish extensive contacts with the hook motif of an adjacent α-subunit (Fig. 1C, 3A). The N-terminal extension of this domain, which is absent in PaMCC (Fig. S5), protrudes into a neighboring β-subunit dimer and includes a short α-helix (residues 66-70) that interacts with the BT-BCCP linker of a second α-subunit (Fig. 3A-B). Residue Y69 in this helix stacks between residues L592 and F596 from the BT-BCCP linker, partially limiting its mobility.

**Figure 3.**
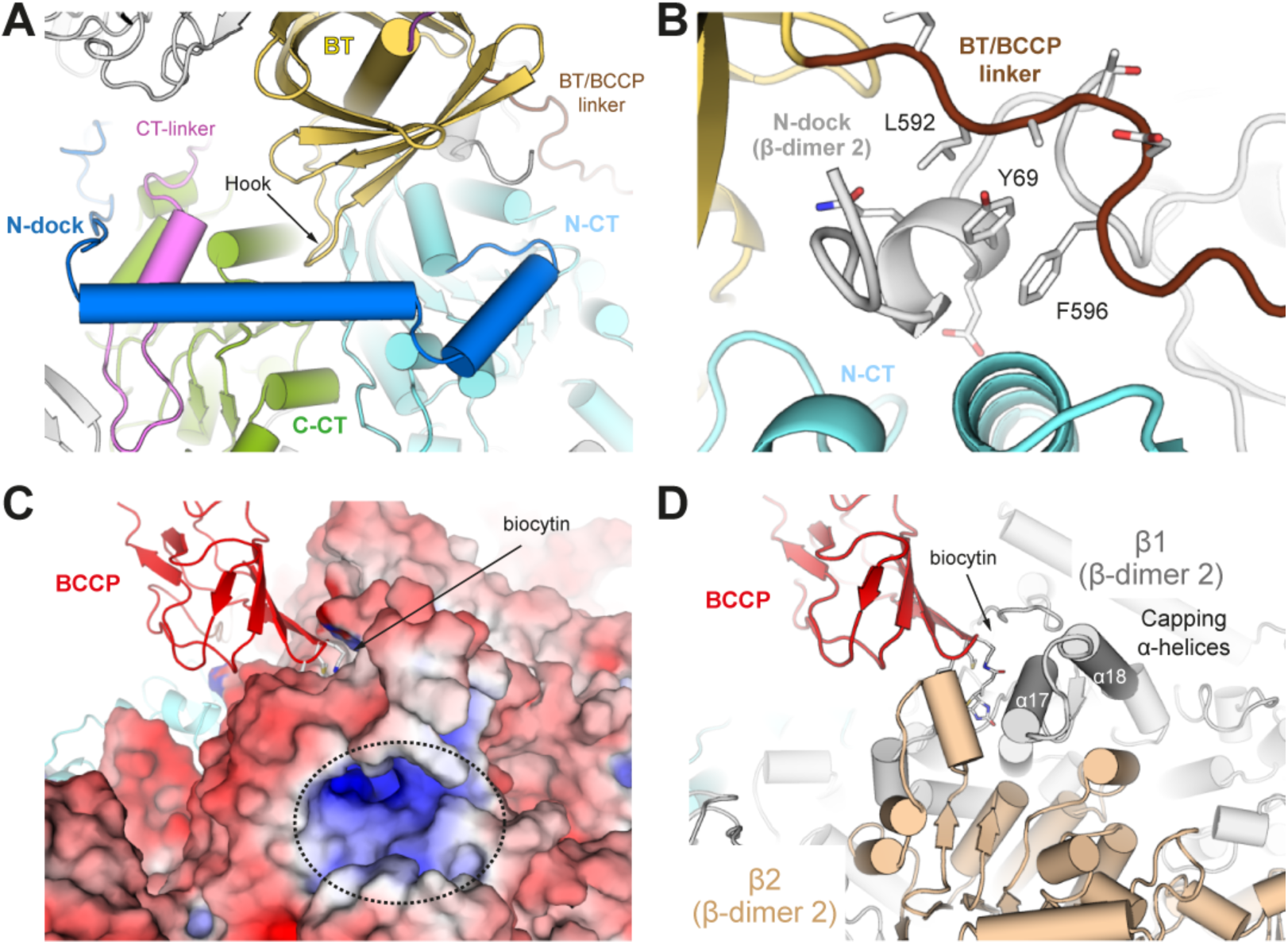
Structural details of β-subunits. (A) Ribbon representation of the interaction region between α- and β-subunits. Note the hook in the BT domain interacting with all domains in the β-subunit. (B) Detailed view of the interaction between the BT-BCCP linker ant the N-dock domain from an aside β-subunit, shown in grey. (C) Electrostatic surface around the β-subunit binding site. A dotted circle indicates the positive region where 3-methylcrotonyl-CoA substrate is expected to bind. (D) Ribbon representation of the β-subunit active site, formed in the interface between two β-subunits, in grey and wheat, showing the capping α-helices that contribute to the substrate-binding pocket.

The N- and C-terminal CT domains (residues 135-348 and 388-601) are connected through the CT-linker (residues 349-387), which is located in the inner region of the MCC barrel next to the closest α-subunit (Fig. 1C, 3A). The active site for 3-methylcrotonyl-CoA carboxylation is placed at the interface between N-CT and C-CT domains belonging to different β-subunits forming each dimer (Fig. 1B-C). Accordingly, these regions exhibit the highest conservation degree in β-subunits (Fig. 1D; Fig. S5). The cryo-EM map shows no density suggesting the presence of 3-methylcrotonyl-CoA in its binding pocket, which harbors a positively-charged narrow tunnel where 3-methylcrotonyl-CoA carboxylation takes place (Fig. 3C), about 80 Å away from the α-subunit active site. Two α-helices (α17 and α18) from the C-CT domain projecting out of the MCC barrel significantly contribute to the formation of the pocket (Fig. 3D).

### Biocytin is protected in a pocket next to the secondary active site

The complete biocytin moiety is clearly defined in the cryo-EM map (Fig. 4A). While covalently linked to the central lysine (K641) in the biotinylation motif of the α-subunit BCCP domain, biotin protrudes from this domain towards the β-subunit dimer located laterally from that interacting with the BT domain of the same α-subunit (Fig. 1B). Biotin, containing a thiophene ring next to an ureido ring, interacts with both β-subunits forming the dimer (Fig. 4B). The thiophene ring is placed in a hydrophobic pocket formed by residue M640 from the BCCP holding the biotin, residues L299 and A303 from the N-CT domain in the distant β-subunit of the dimer, and residues I432 and V466 from the C-CT domain in the proximal β-subunit of the dimer. Additionally, the ureido ring interacts with the main chain of residues T462 and F464 and the side chain of residue Q534 in the latter domain. All these residues, including those interacting with biotin through the main chain, are conserved (Fig. S4 & S5). The oxygen atom within this biotin moiety is directed to a narrow tunnel connecting the biotin binding site with the binding site for 3-methylcrotonyl-CoA (Fig. 4C). In this unanticipated location, no density is observed for the carboxyl moiety of biotin. Our results suggests that, upon 3-methylcrotonyl-CoA carboxylation and product release, the BCCP domain is retained next to the β-subunit active site, with biotin protected in an alternative pocket next to that required for carboxylation.

**Figure 4.**
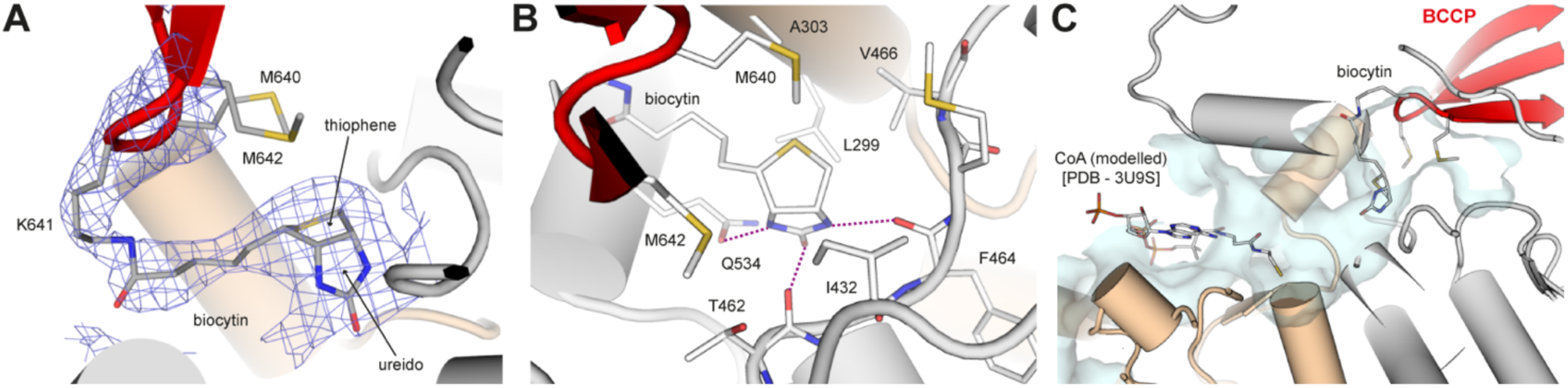
Structural details of the alternative biotin-binding pocket. (A) Cryo-EM map around biocytin. (B) Biocytin interactions inside the binding pocket. Note that residues from both β-subunits, in grey and wheat, are involved in the interaction. (C) Tunnel connecting the alternative biotin-binding pocket and the substrate binding site. For clarity, CoA from the PaMCC structure (PDB - 3U9S) is shown as sticks.

### Dynamics of the BC domain shows a coordinate movement of α-trimers

The largest inter-chain interaction is established between β-subunits forming dimers (3504 Å^2^ area), which are placed tail-to-tail in the central region of the MCC barrel (Fig. 1B). A total of 24 hydrogen-bonds (H-bonds) and 4 salt bridges are observed in this interface (Table S2). The interface between β-subunits forming each trimeric ring is three times smaller (1356 Å^2^ area) where 17 H-bonds and 5 salt bridges are observed (Table S2). These observations indicate that the β-subunit dimer is the primary building block for the assembly of the MCC barrel core. In contrast, the buried surface connecting adjacent α-subunits is smaller (494 Å^2^) (Fig. 1B), suggesting that α-trimers could be more flexible than the β-core. This may also indicate that α-subunits assemble on the MCC barrel at a later stage, as shown for the PCC enzyme (Lee et al. 2023), supported by observation of pre-formed β-hexamers particles with only one α-trimer in negative-stain grids (Fig. S1).

The main interacting region between α- and β-subunits is established between the BT domain of the former and residues from all domains in the latter (1100 Å^2^). A total of 10 H-bonds and 1 salt bridge are established (Table S2), most of them between the BT hook of α-subunits and the N-terminal region (residues 137-155) of the N-CT domain in β-subunits (Fig. 3A). Beside this principal contact with the closest β-subunit, each α-subunit interacts with the β-subunit located aside from the closest one (Fig. 1B). First, the N-terminal region (residues 64-78) in the N-dock domain of the aside β-subunit interacts with the BT domain and the BT-BCCP linker. Second, the BCCP domain interacts with the N-CT domain of the aside β-subunit (874 Å^2^). These lateral contacts are important for BCCP activities during MCC catalysis, as they partially limit BCCP swinging while allowing alternative binding configurations next to the β-subunit active site (see below).

To study the dynamics the MCC enzymatic complex, we analyzed the conformational distribution of the particles in our cryo-EM dataset. We observed that the β-subunit core is quite rigid and only minor rearrangements are observed (Movie 1). In contrast, α-subunits are highly dynamic, especially the BC domains at the edges of the MCC barrel. On each side of the barrel, BC domain trimers pivot over their corresponding BT domains to approach the β-subunit core on one side of the barrel. Unexpectedly, the three BC domains forming each trimer move together, such that the trimeric interaction is preserved. Consequently, when one BC domain approaches the β-subunit core, the other two BC domains in the trimer travel together, thus moving away from the core. This suggests that only one of the three α-subunit active sites can approach the β-subunit core at a time, then travels back to allow the approach of α-subunit another active site towards the core. Interestingly, the two α-subunit trimers appear to pivot coordinately so that both trimers approach the β-subunit core from the same side (Movie 1).

### The TbMMC structure represents a resting state

Superposition of the TbMCC structure with those of PaMCC and LtMCC shows an overall similarity, with root-mean-square deviations (RMSD) of 0.7 and 1.2 Å, respectively. Likely due to the filamentous nature of the structure, LtMCC presents a more extended barrel with a height of 212 Å, while PaMCC and TbMCC barrels are 201 Å high. The most prominent differences concern α-subunits, which in LtMCC are less tightly packed against β-subunits (Fig. 5A), further supporting the flexible nature of the α/β interface. BC and BT domains in α-subunits of LtMCC shift away from β-subunits by 4.5 and 3 Å, respectively, compared to their position in TbMCC. As a result, the contact surface between α and β-subunits is smaller in LtMCC than in TbMCC, i.e. 777 versus 1096 Å^2^. In contrast, PaMCC presents a larger interaction surface between α- and β-subunits (1492 Å^2^), mainly due to an extended hook motif in the α-subunit (Fig. S4). The extended PaMCC hook partially overlaps with β-subunit residues at the interface between the N-CT and C-CT domains, including the CT-linker, in the structures of MCCs from eukaryotic parasites. Notably, β-subunit residue Q592 in TbMCC forms an H-bond with R183 that reinforces the interaction between the N-CT and C-CT domains, while the equivalent residues in PaMCC interact with the α-subunit hook (Fig. 5B). This indicates a tighter packing of β-subunits at the expense of a weaker interaction between α- and β-subunits in parasite MCCs.

**Figure 5.**
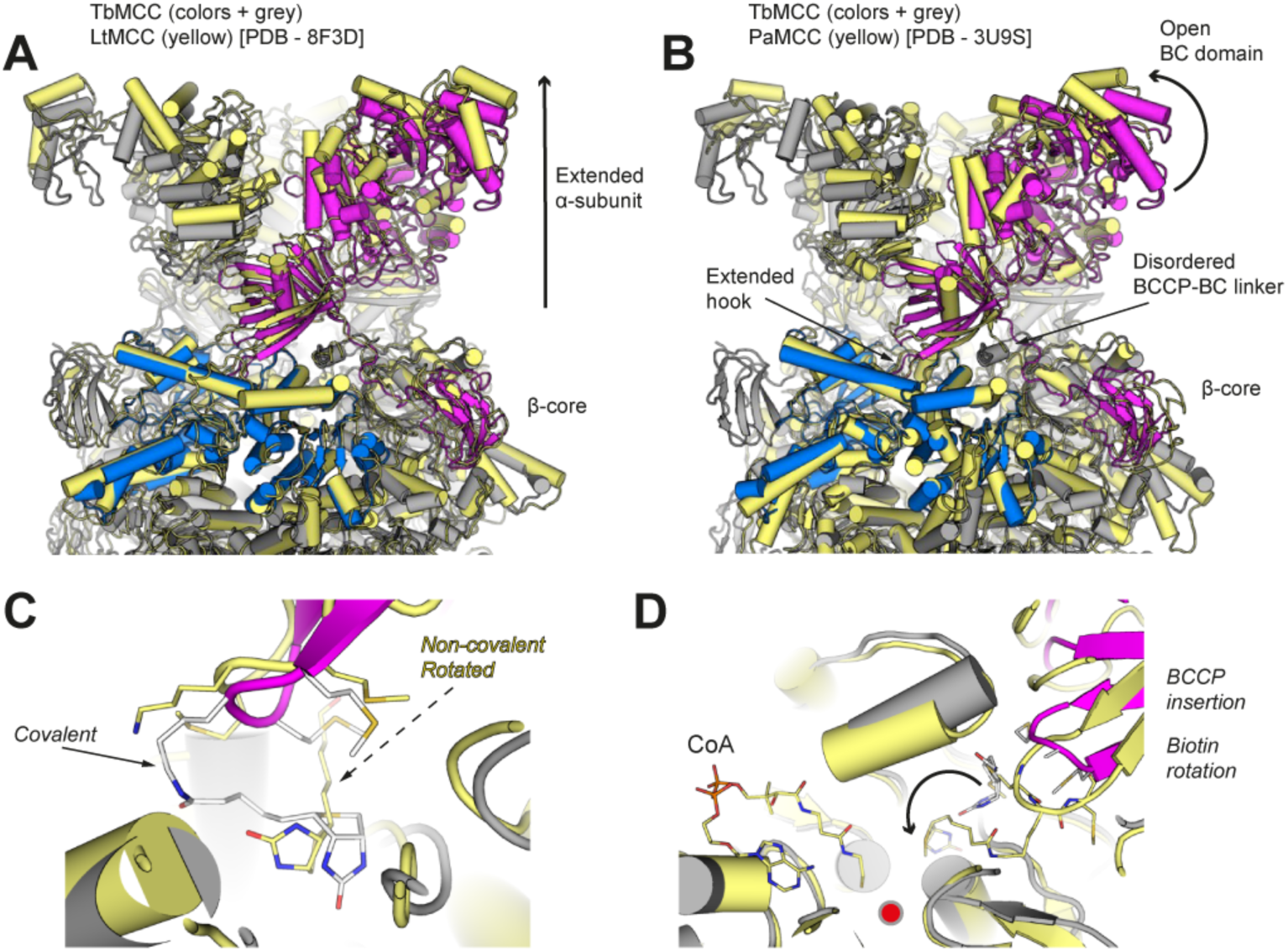
Structural comparison of TbMCC with reported MCCs. (A) Structural comparison with the cryo-EM structure of LtMCC filaments (PDB - 8F3D), in yellow, showing the displacement of α-trimers due to filament formation. (B) Structural comparison with the crystal structure of PaMCC (PDB - 3U9S), in yellow, where an open BC domain was described and is involved in crystal contacts. (C) Detailed view of the β-subunit active site, compared with that of LtMCC in yellow. Note that, in the latter, biotin is not attached covalently to the lysine residue in the biotinylation motif. (D) Detailed view of the β-subunit active site, compared with that of PaMCC in yellow, where the BCCP domain and biotin are more deeply inserted into the active site, for catalysis.

The α-subunit active site in TbMCC is narrower than in the structures of previously-reported MCCs (Fig. 5A-B). In opposition to the already discussed *E. coli* BC structures (Thoden et al. 2000), these open configurations are likely due to contact with neighboring dodecameric barrels in LtMCC filaments or PaMCC crystals, both involving large interaction areas in this region in the absence of substrates (Fig. 2C). Regarding the β-subunit active site, the structure of PaMCC complexed to CoA shows that the biocytin moiety is placed deeper in the active site, next to the substrate, while biotin in LtMCC is not covalently attached to the protein (Fig. 5C-D). This correlates with closure of this active site in PaMCC. Interestingly, the BT-BCCP linker in the α-subunit is disordered in PaMCC, in contrast to parasite MCCs showing extensive interactions between this linker and the N-terminal extension in the N-dock domain of a neighbor β-subunit (Fig. 3B), which is unique in eukaryotes (Fig. S4). The interaction between the N-dock N-terminal extension and the BCCP-BC linker is stronger in TbMCC as compared to LtMCC. Altogether, these observations suggest that the PaMCC structure likely mimics a post-catalytic state, while that of TbMCC represents the resting state of the enzyme.

## Discussion

The structure of MCC from *T. brucei* provides new details that deepen our understanding of this mitochondrial enzyme, essential for leucine metabolism. While the core of its cylindrical structure, composed by six copies of the β-subunit, superposes well with already known structures from *P. aeruginosa* and *L. tarentolae*, the outer regions of the cylinder exhibit interesting details. Notably these regions, composed by α-subunit trimers, are now observed in their native state due to lack of interactions arising from crystal packing (as for PaMCC) or filament formation (as for LtMCC). As a result, the flexibility of the two α-trimers can be analyzed.

While the A- and C-motif of the BC domain superpose with *E. coli* carboxylase, the B-motif presents an intermediate state between open and closed conformations, with a sulphate ion in the bicarbonate binding site of this highly-conserved region (Fig. 2). As seen in carboxylases, binding of Mg^2+^/ATP induces this flexible B-motif to close the active site, which is widely open in PaMCC and LtMCC. Unexpectedly, dynamic analysis of our cryo-EM data shows that not only the B-motif is flexible, but each entire α-trimer is able to pivot so that one of the three BC domains in the trimer approaches the β-core (Movie 1). While this movement is limited, this result confirms that α-trimers can reduce the distance that the BCCP domain needs to cover in order to reach the α-subunit active site. Nevertheless, the BT-BCCP linker in TbMCC is partly fixed due to contacts with the N-dock domain of a neighboring β-subunit (Fig. 3), as also observed for LtMCC. Residues in the BT-BCCP linker contacting the N-dock domain are absent in PaMCC or mammalian MCCs (Fig. S4), suggesting that only parasite MCCs partly restrict BCCP swinging. Interestingly, PaMCC and mammalian MCCs present hook regions that are longer than those in parasite MCCs. As seen in the structure of PaMCC, this involves a larger interaction area with β-subunits, which might restrict flexibility of α-trimers in species with longer hooks.

Biotin in TbMCC is covalently bound to lysine in the BCCP domain, otherwise than seen in LtMCC (Fig. 5). However, as opposed to PaMCC in the presence of the substrate analog CoA, biocytin in TbMCC locates more superficially to the β-subunit active site, occupying a secondary pocket that connects to the active site through a narrow channel (Fig. 4). This resting position for the BCCP domain may protect it from degradation and/or hamper undesired biotin interactions while MCC activity is not taking place. The structure of the secondary biotin-binding pocket may assist drug discovery against trypanosome infections. Moreover, as the residues in the pocket are conserved (Fig. S5), it may also assist the search for inhibitors of human MCC that could refrain tumor progression.

Based on available structures and that reported here, a general model for MCCs can be proposed (Fig. 6). As stated for LtMCC, the filament state corresponds to an inactive configuration, due to the intra-filament interactions that immobilize BC domains. MCC in this quiescent state in the mitochondrial matrix can be easily reactivated when required. Free dodecameric complexes, as in the TbMCC structure, represent a resting state in the reaction where biotin, already bound to lysine in the biotinylation motif, is protected next to the β-subunit active site but ready to be translocated to the α-subunit active site for carboxylation. The coordinated movement of α-trimers observed in TbMCC suggest that both BC and BCCP domains could move for biotin carboxylation, as stated for the dual-swinging model. According to our results, BCCP swinging can cover most of the distance between active sites, while α-trimer pivoting is more limited. α-trimer pivoting likely regulates sequential carboxylation of the three BCCP domains, as only one of the trimers can approach the barrel core at a time. Upon biotin carboxylation at the α-subunit active site, the BCCP can travel back to the interface between β-subunits for the second step in the enzymatic reaction. While this second step takes place at one β-dimer, α-trimer pivoting would allow concomitant carboxylation of a second biotin moiety at a neighboring α-subunit active site. The PaMCC structure, with CoA mimicking the carboxyl acceptor and carboxyl-free biotin, represents the post-reaction state for the second step. Upon this reaction, MCC may continue its enzymatic task, fall back in the resting state, or be stored in the inactive filamentous state. Additional structural analysis in the presence of substrates or analogs will shed further light on the mechanisms of MCC activity.

**Figure 6.**
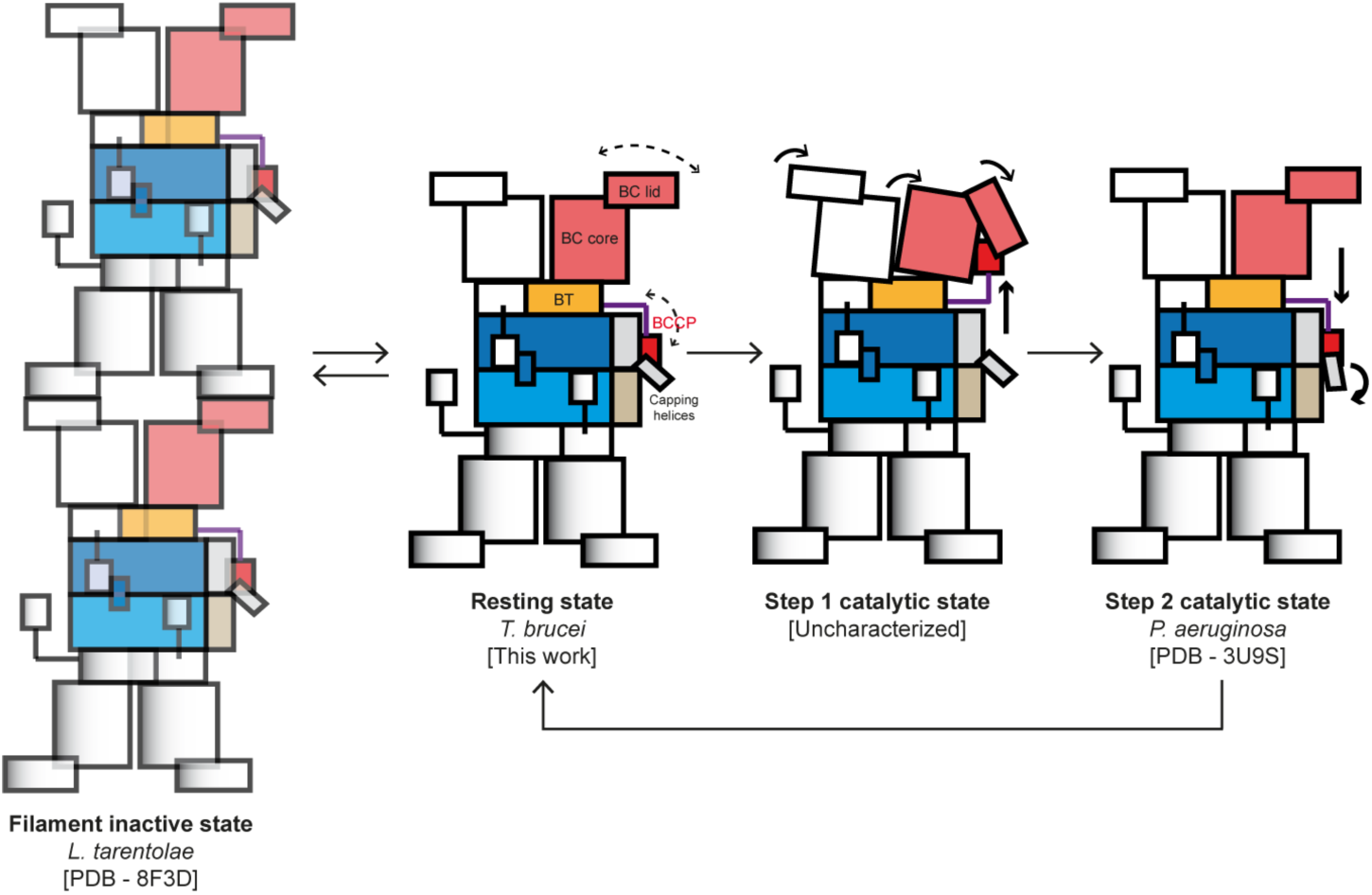
Mechanistic model for MCC catalysis. Schematic representation of the general activity of the MCC enzyme, showing the different states represented by the three known MCC structures and a putative state for the first catalytic reaction, currently uncharacterized structurally. β-subunits are in blue, while α-subunits are in white or grey. Only one α-subunit that is colored in salmon, yellow, purple and red for BC domain, BT domain, CT-BCCP linker and BCCP domain, respectively. Flexible regions in the TbMCC structure are shown as dotted arrows, while thick arrows indicate domain movements for the catalytic mechanism.

## Materials and methods

### Purification of 3-methylcrotonyl-CoA carboxylase from *T. brucei*

Harvested *T. brucei* cells were resuspended in lysis buffer (25 mM Hepes pH 7.5, 150 mM ammonium sulphate, 5 mM MgCl2, 5% Glycerol, 0.5% Triton X100, 2 mM BME and one tablet of EDTA-free protease inhibitors (Roche)), and lysed by sonication. The lysate was clarified by centrifugation at 35 Krpm for 30 min in a 45ti rotor (Beckman). The supernatant was filtered (0.45 um) and loaded in a Strep-tactin column (1 ml, IBA) equilibrated with buffer containing 25 mM Hepes pH 7.5, 150 mM Amm. Sulf., 5 mM MgCl2, 5% Glycerol and 2 mM BME. After washing the column with 10 ml of the same buffer the sample was eluted with 3 ml of 25 mM Hepes pH 7.5, 150 mM Amm. Sulf., 5 mM MgCl2, 5% Glycerol, 2 mM BME and 50 mM Biotine. Sample was concentrated using a centrifugal filter device (Amicon Ulltra-05 10K), plunged in liquid nitrogen and stored at -80 C for grids preparation.

### Negative stain electron microscopy

Electron micrographs were recorded on a JEOL 1230 electron microscope operated at 100 kV. 25 images were collected using a CMOS TVIPS TemCam-F416 detector under the control of the EM-Tools software (TVIPs). Data processing was done using Relion 3.1. A total of 37313 particles were picked. After several rounds of 2D classification 28348 particles were finally selected for 3D refinement. The final 3D volume reached a resolution of 17 A. The PaMCC structure (Huang et al. 2012) was fitted in this map using Chimera (Pettersen et al. 2004).

### Identification of proteins by MALDI-TOF-TOF peptide mass fingerprinting

Individual SDS-PAGE protein bands were cut manually and digested with trypsin (Thermo). Samples were analyzed with an Autoflex III TOF/TOF mass spectrometer (Bruker-Daltonics).Typically, 1000 scans for peptide mass fingerprinting (PMF) and 2000 scans for MS/MS were collected. Automated analysis of mass data was performed using FlexAnalysis software version 3.4 (Bruker-Daltonics). MALDI–MS and MS/MS data were combined through the BioTools 3.2 program (Bruker-Daltonics) to SwissProt 2021_02 database using MASCOT software 2.6 (Matrix Science). Relevant search parameters were set as follows: enzyme, trypsin; fixed modifications, carbamidomethyl (C); oxidation (M); 1 missed cleavage allowed; peptide tolerance, 50 ppm; MS/MS tolerance, 0.5 Da. Protein scores greater than 75 were considered significant (p<0.05).

### Cryo-EM sample preparation

C-flat copper 300 mesh 1.2/1.3 holey carbon grids (Protochips) covered with a home-made continuous carbon film were glow discharged for 20 s and a current of 25 mA using a Glow Discharge device Quorum GloQube dual camera. Then 3 µl of sample at 0.07 mg/ml concentration were applied to the glow-discharged grids and blotted for 2 s with blotting force -5 using a FEI Vitrobot (Thermo Fisher Scientific) at a temperature 10 °C and 100% humidity. Grids were vitrified by plunging into liquid ethane cooled with liquid nitrogen, and stored in liquid nitrogen for later imaging.

### CryoEM image acquisition

Movies were acquired in a FEI Titan Krios (ThermoFisher) electron microscope working at an acceleration voltage of 300 keV. A total of 4530 movies (40 frames each) were collected with a K3 summit (Gatan) direct electron detector operated in ‘super-resolution’ mode, using the EPU software, with a defocus value range from -1 to -3 µm, a pixel size of 0.8238 Å/pixel, a dose rate of 18.361 e–/px/s and a total dose of 38.3 e/Å^2^.

### CryoEM image processing

Movies were aligned and motion corrected with MotionCor2 (Zheng et al. 2017) and subsequent processing was performed in CryoSPARC v4.2 (Punjani et al. 2017). Patch CTF estimation was followed by visual inspection, and 157 movies were rejected using the ‘curate exposures’ job. From the remanent 4377 micrographs, initial picking was done with ‘blob picker’ using an ellipse template of 200 by 290 Å. These particles were 2D classified and the resulting 2D averages were used as templates to pick a total of 490k particles. After extraction with a 540 pixel box, a reference-free 2D classification was performed and the best classes, including 188k particles, were selected. Ab-initio reconstruction was used to generate an initial 3D model, which was refined applying C3 symmetry, also used in subsequent 3D refinements. To further clean the dataset, a heterogeneous refinement was performed using both the initial 3D model and a reference generated from discarded 2D classes as a decoy volume. This resulted in a final set of 126k particles that, after homogeneous refinement, yielded a map at 2.7 Å resolution. These particles underwent iterative rounds of local defocus refinement and CTF aberration refinement (tilt, trefoil, spherical aberration and tetrafoil). The final reconstruction was obtained by non-homogeneous refinement, yielding a map with a global resolution of 2.5 Å. Local resolution was calculated and 3DFSC (ref) was used to asses directional FSC.

The ‘3D Flex’ job (Punjani and Fleet 2023) in CryoSPARC was used to analyze the conformational distribution of the sample. Particles were cropped to a box size of 320 pixels and then downsampled to 160 pixels. A Flex Mesh was prepared with 20 tetra cells and no segmentation. Then the 3D Flex model was trained with 64 hidden units and a rigidity of 2. While several numbers of latent dimensions were tried, the series of volumes generated showed an indistinguishable movement, so only the first dimension was considered. The use of a segmented mesh showed only the movement of one monomer, likely due to inability to properly set the sub-mesh connections.

### Model building and refinement

An AlphaFold2 prediction (Jumper et al. 2021) was used as first model for both chains. Initial model placement into the cryo-EM map was done using UCSF Chimera (Pettersen et al. 2004). The map was cut out around a asymmetric αββα unit, using Phenix (Liebschner et al. 2019). Manual modification and model building was done using Coot (Emsley et al. 2010). Real space refinement using secondary structure restraints was done using Phenix. Finally the model was triplicated according to the C3 symmetry and fitted in the full EM map for the last rounds of refinement. Contact surface properties were calculated using the PISA Web server (Krissinel and Henrick 2007). Protein volumes were calculated using the CASTp web server (Tian et al. 2018). Figures were prepared with Pymol (ref) and ChimeraX (ref).

## Acknowledgments

The identification of proteins by MALDI-TOF-TOF peptide mass fingerprinting was carried out at the Proteomics and Genomics Facility (CIB-CSIC, Madrid, Spain), a member of ProteoRed-ISCIII network. Grid preparation and negative stain electron microscopy experiments were done at the Electron Microscopy Facility (CIB-CSIC, Madrid, Spain). We thank David Gil-Carton at BREM Instituto Biofisika (Leioa, Spain) for support with cryo-EM data collection.

## Data availability

The cryo-EM map and the derived atomic model have been deposited in the Electron Microscopy Database and Protein Data Banks, respectively, under accession codes EMD-xxxx and PDB-yyyy.

